# Identification of a primitive intestinal transcription factor network shared between oesophageal adenocarcinoma and its pre-cancerous precursor state

**DOI:** 10.1101/373068

**Authors:** Connor Rogerson, Edward Britton, Sarah Withey, Neil Hanley, Yeng S. Ang, Andrew D. Sharrocks

## Abstract

Oesophageal adenocarcinoma (OAC) is one of the most frequent causes of cancer deaths and yet compared to other common cancers, we know relatively little about the molecular composition of this tumour type. To further our understanding of this cancer we have used open chromatin profiling to decipher the transcriptional regulatory networks that are operational in OAC. We have uncovered a transcription factor network that is usually found in primitive intestinal cells during embryonic development, centred on HNF4A and GATA6. These transcription factors work together to control the OAC transcriptome. Importantly, we show that this network is activated in Barrett’s oesophagus, the putative precursor state to OAC thereby providing novel molecular evidence in support of stepwise malignant transition. Furthermore, we show that HNF4A alone, is sufficient to drive chromatin opening and activation of a Barrett’s-like chromatin signature when expressed in normal human epithelial cells. Collectively, these data provide a new way to categorise OAC at a genome scale and implicate HNF4A activation as a potential pivotal event in regulating its malignant transition from healthy cells.

## Introduction

Oesophageal adenocarcinoma (OAC) is one of the eight most common cancers in the Western world and yet it has very low survival rates (Pennathur et al., 2013). One reason for the poor prognosis is the lack of tailored therapies due to the relative paucity of molecular knowledge compared to other cancers. We are beginning to understand the molecular mechanisms underpinning this disease, chiefly through genomic sequencing studies, which have revealed multiple genes that are recurrently mutated in OAC (Dulak et al. 2013; Weaver et al., 2014; Ross-Innes et al., 2015; Frankell et al., 2018). However, with the exception of *TP53*, the overall incidence of mutations in individual genes is low. By grouping genes in broader functional categories, frequently mutated pathways have been uncovered, such as chromatin remodelling complexes, the RAS-ERK signalling pathway and cell cycle control pathways. Another broader category of interest is comprised of genes encoding transcriptional regulatory proteins which include both GATA4 and GATA6. Together amplification of the genes encoding these two transcription factors has been reported in up to 40% of OAC cases (Lin et al., 2012; The Cancer Genome Atlas Network, 2017). This suggests that transcriptional network rewiring might be an important element in the progression of OAC.

There are persuasive arguments in favour of OAC developing from a pre-existing condition known as Barrett’s oesophagus (Desai et al., 2012; Burke and Tosh, 2012). OAC and Barrett’s share a large number of molecular markers. For example, many of the mutations found in OAC are already present in Barrett’s, including in *TP53*, suggesting that they may help drive the transition to Barrett’s as a stepwise process to OAC, rather than to OAC directly from healthy tissue (Stachler et al., 2015; Ross-Innes et al., 2015). In contrast, focal gene amplifications tend to arise following the transition from Barrett’s to OAC suggesting that these may play a more important role in establishing the OAC state (Lin et al., 2012; Ross-Innes et al., 2015; Yamamoto et al., 2016). Morphologically, the normal oesophagus consists of a stratified squamous epithelium, but Barrett’s differs significantly from this and instead resembles a columnar epithelium, typically found in the more distal gastrointestinal tract (reviewed in Spechler and Souza, 2014). Several models have been proposed for how this metaplastic transition occurs including transdifferentiation from normal oesophageal epithelial cells (Stairs et al., 2008; Vega et al., 2014), colonisation by migrating cells of gastric origin (Quante et al., 2012) or more recently, changes to cells from the gastroesophageal junction (GOJ) (Jiang et al., 2017).

Despite these advances, mutational signatures have not yet provided a unified insight into how OAC is initiated and maintained. Changes to the epigenetic landscape might be a major contributing factor, and this scenario has been implicated in other cancers (Davie et al., 2015; Rendeiro et al., 2016). To gain insights into the molecular mechanisms that are operational in OAC, we therefore turned to ATAC-seq to study the open chromatin landscape as this has been successfully applied to studying cell fate transitions such as neuron differentiation from fibroblasts (Wapinski et al., 2017), haematopoiesis (Yu et al., 2016), epidermal differentiation (Bao et al., 2015) and embryonic development (Cusanovich et al. 2018). We recently applied this approach on cell line models of OAC to uncover AP1 as a major transcriptional regulator in this context (Britton et al., 2017). In the current study, we interrogated the chromatin landscape of patient samples, and uncovered a complex regulatory network in OAC, shared with Barrett’s oesophagus, consisting of several transcription factors that usually operate during early human intestinal development (HNF4A, GATA6, FOXA1 and HNF1B). Among these, HNF4A was demonstrated to possess pioneering activity and is sufficient to drive chromatin opening and induce a Barrett’s-like phenotype.

## Results

### Identification of a network of transcription factors active in OAC

Previously we used ATAC-seq to profile the open chromatin landscape of OAC cell lines and identified AP1 as an important transcription factor family in controlling the transcriptional networks in OAC (Britton et al., 2017). To further interrogate the transcriptional networks operating in OAC, we decided to take an alternative approach, starting with clearly defined open chromatin datasets from OAC patient samples rather than a diverse set of OAC-derived cell lines. We previously validated our cell line-derived results by profiling the open chromatin of six patient-derived biopsies but this revealed two distinct subclusters of OAC samples based on their open chromatin landscapes; one that clustered with normal samples and one that was unique (Britton et al., 2017). We revisited this issue by performing subclustering by PCA analysis following the addition of an additional paired normal and OAC dataset and again we observed a clear partitioning of samples (Supplementary Fig. S1A). We therefore re-focussed our attention on the four OAC samples which are clearly distinct from the normal samples (T_002, T_003, T_005, T_006; Supplementary Fig. S1A).

To derive a unified dataset of open chromatin regions in oesophageal-derived tissue, we combined all the ATAC-seq data from three normal oesophageal tissue samples (matched with T_003, T_005 and T_006) and these four OAC samples and recalled the open accessible chromatin regions from this combined data. We selected the top 50,000 open regions (Supplementary Table S1A) and found that these were roughly equally divided between promoter-proximal, intragenic and intergenic regions (Fig. 1A). PCA analysis based on these peaks confirmed the distinct clustering of the normal and OAC samples (Fig. 1B). Next we identified regions that are differentially accessible between the normal and OAC samples, by comparing the average signals in each type of sample. This yielded a total of 1,438 differentially accessible regions, the vast majority of which (95%) are located in intra- or intergenic regions (Fig. 1C and D; Supplementary Table S1B and S1C). Clustering analysis using these differentially accessible regions clearly separates the normal and OAC samples and revealed that chromatin opening is more prevalent than closing in OAC (71% regions are more open in OAC) (Fig. 1C and D). GO terms analysis revealed that these open regions are associated with genes with functions such as “cell development”, “response to hormone stimulus” and “response to wounding” (Supplementary Fig. S1B). We then asked whether opening of chromatin corresponds with an increase in gene expression. We interrogated RNAseq data from normal oesophageal and OAC tissue (Maag et al., 2017) and found that genes associated with a region demonstrating an increase in accessibility (promoter and non-promoter) in OAC also show elevated levels of expression in OAC (Fig. 1E).

**Figure 1.**
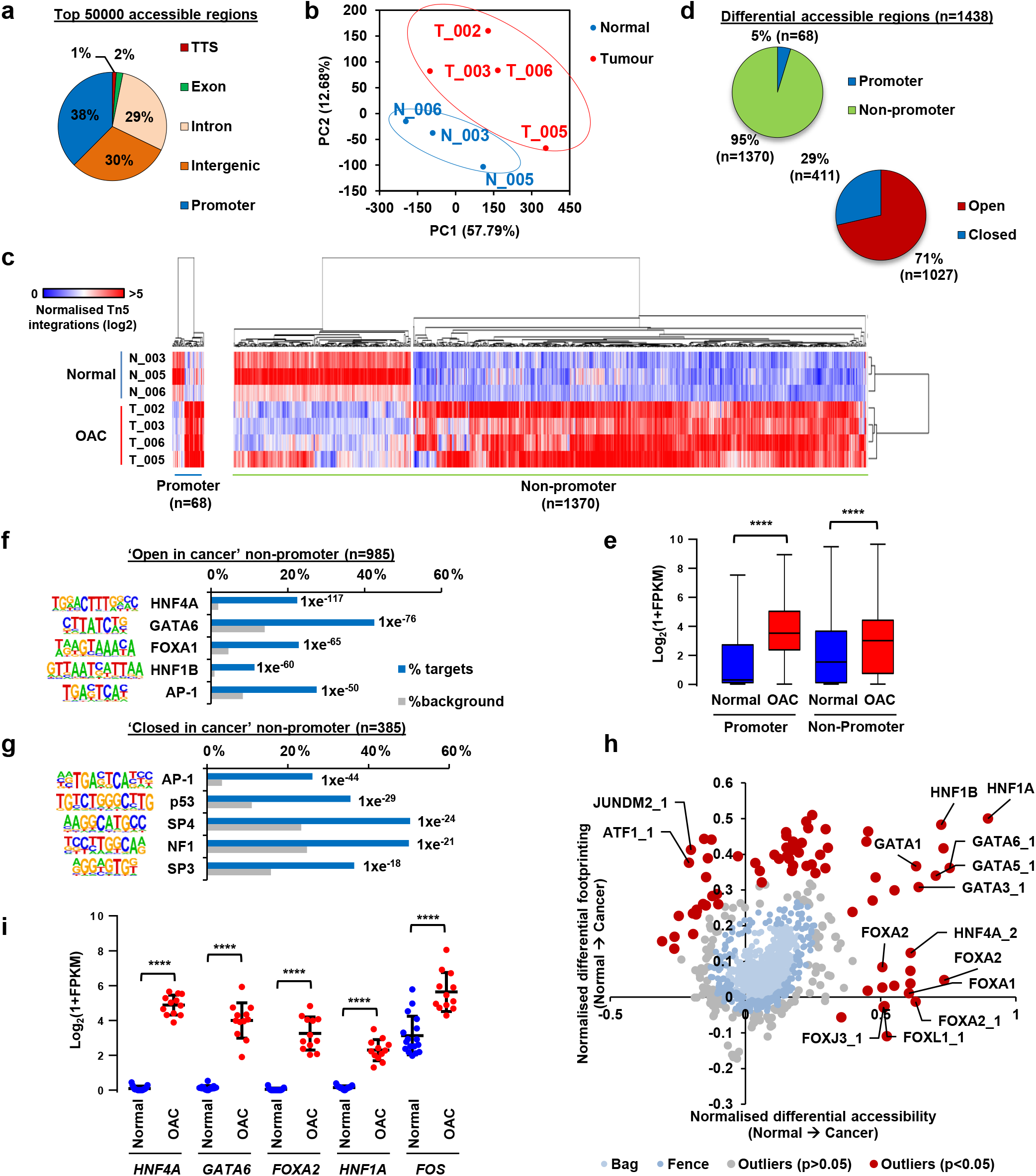
Open chromatin profiling reveals dynamic chromatin accessibility in OAC. (A) Genomic distribution of the top 50,000 significant open chromatin regions in combined ATAC-seq data from normal and tumour tissue. “Promoter” refers to −2.5 kb to +0.5 kb relative to the TSS. (B) PCA plot of ATAC-seq signal across the top 50,000 ATAC-seq regions in 3 normal tissue samples (blue) and 4 tumour tissue samples (red). (C) Heatmap of normalised Tn5 cleavage events in a +/− 250 bp region surrounding the summits of differentially accessible promoter and non-promoter regions (linear fivefold difference, p<0.05). Hierarchical clustering was performed on samples and regions using 1-Pearson’s correlation. (D) Pie charts representing the proportion of differentially accessible regions that are located in promoter and non-promoter regions (left) and regions that are five-fold more open or closed in tumour tissue (right). (E) Box plot of Log_2_ 1+FPKM values of genes associated with promoter and non-promoter regions that show increased accessibility in OAC (red) compared to normal tissue (blue). Whiskers represent 1.5 × IQR. (F and G) Top five DNA motifs derived from *de novo* motif discovery and their associated transcription factor that are enriched in ‘open in cancer’ (F) or ‘closed in cancer’ (G) non-promoter regions. The frequency of motif occurrence is shown and the motifs are sorted by p-value. (H) Scatter bag plot of differential chromatin accessibility (x-axis) and footprinting (y-axis) depth around human transcription factor binding motifs in normal and cancer tissue. Significant outliers (p<0.05) are represented in red. Transcription factor motifs with enrichment in ‘Open in Cancer’ are labelled. (I) Plot of Log_2_ FPKM values of transcription factors with enriched motifs in ‘open in cancer’ regions in RNAseq data from normal (blue) or OAC (red) tissue (Maag et al., 2017). Mean is represented by black bar with standard deviation shown above and below. **** represents p<0.0001.

To gain further insights into the regulatory networks that act on these differentially open chromatin regions, we first identified DNA motifs that are over-represented in the differentially accessible non-promoter regions (Fig. 1F; Supplementary Table S2A). As expected from our previous work, we found AP1 motifs in the differentially open regions in OAC but additionally we found greater enrichment of binding motifs for the HNF4, GATA, FOXA and HNF1 subfamilies of transcription factors (Fig. 1F). Of these motifs, HNF4 was also found at promoter regions (Supplementary Fig. S1C). In contrast, with the exception of AP1, a different set of DNA motifs are associated with regions closed in cancer, with the p53 motif being the most prevalent (Fig. 1G). In addition to demarcating open chromatin, ATAC-seq can also reveal the precise sequence bound by a transcription factor and thus protected from digestion (the ‘footprint’) and subsequently the underlying DNA sequence motifs. For robustness, we used two complementary approaches. By the Wellington algorithm (Piper et al., 2013), we again identified the motifs for HNF4A/G and GATA6 (Supplementary Fig. S1D). Next we used BaGFoot (Baek et al., 2017) to identify motifs that showed evidence of changes of occupancy through altered “footprinting” in the OAC samples across all of the top 50,000 open regions in the tissue-derived ATAC-seq datasets. Again, we identified motifs for the HNF4, GATA, FOXA and HNF1 transcription factor subfamilies in this analysis which all exhibited greater localised accessibility in cancer, with the GATA6 and HNF1B motifs showing particularly strong increases in footprinting depth across the motif itself (Fig. 1H; Supplementary Fig. S2). Conversely, the p53 motif showed evidence of reduced footprinting depth and local chromatin accessibility (Supplementary Fig. S2). Given the enrichment of these TF motifs, we next determined whether any of the transcription factors which recognise these motifs are upregulated in OAC. Again, we used RNAseq data from normal oesophageal and OAC tissue (Maag et al. 2017) and found that GATA4/6, and HNF4, FOXA and HNF1 family members are all upregulated in OAC samples, albeit with different subfamily members in individual patients (Fig. 1I; Supplementary Fig. S1E). Furthermore, representative genes encoding transcription factors from these subclasses also show increased ATAC-seq signal in their putative regulatory regions in OAC tissue compared to normal tissue (Supplementary Fig. S1F), consistent with their transcriptional upregulation. To determine whether these transcription factors may form a network in OAC, we focussed on HNF4A and GATA6 motifs in regions of open chromatin and counted the frequency of co-occurring motifs for HNF4A, GATA6, FOXA1, HNF1B and AP1 within the same regions. For both HNF4A and GATA6 motifs, there is a significantly different distribution of co-occurring motifs in regions that are more accessible in OAC compared to randomly selected genomic regions containing either of these motifs (Supplementary Fig.1G). This is particularly marked for the co-occurrence of HNF4A with GATA6 motifs and suggests the existence of a complex transcription factor network in OAC.

Given the conflicting theories about how OAC arises, we wanted to explore where else this regulatory pattern of transcription factor activity might be observed. Given the highest enrichment for the GO term ‘regulation of cell development’ (Fig. S1B), we hypothesized that OAC might be mimicking aspects of human embryogenesis, and in particular distal foregut endoderm (Gerrard et al, 2016). Indeed, the cohort of nine transcription factors were most highly enriched in embryonic derivatives of distal foregut endoderm (stomach and liver) compared to proximal derivatives (thyroid or lung) or tissues mainly derived from other germ layers [e.g. brain (ectoderm) or heart ventricle, adrenal or kidney (mesoderm)] (Supplementary Fig. S3). However, strong co-expression was not observed in any single tissue type. To explore this further, we integrated the human embryonic data with RNAseq from differentiating human pluripotent stem cells. The strongest enrichment was observed at the stage of foregut endoderm/early liver differentiation, where representative members from each of the four subfamilies are co-expressed.

Collectively, these data implicate HNF4A, GATA6, FOXA2/3 and HNF1B as a group of transcription factors specific to OAC controlling gene expression predominantly through distal regulatory sites. This complement of factors points to OAC arising due to a reactivation of a distal human embryonic foregut phenotype.

### Identifying the GATA6 and HNF4A cistrome

As GATA6 and HNF4A show the highest motif occurrence and the highest differential expression in OAC, we focussed on these two transcription factors, and aimed to determine their role in controlling gene expression in OAC cells. We first sought a suitable cell line whose open chromatin environment resembled that found in the patient-derived OAC samples. Initially we used PCA to cluster the open chromatin regions from four OAC-derived cell lines with the patient-derived samples. OE19 cells were clearly identified as most closely resembling the open chromatin landscape of primary OAC (Supplementary Fig. S4A). This was confirmed by Pearson’s correlation analysis (Supplementary Fig. S4B). Additionally, both GATA6 and HNF4A levels are elevated in OE19 cells compared to the other cell lines (Supplementary Fig. S4C). *CLRN3* has been identified as a HNF4A transcriptional target in iPSC-derived hepatocytes (Mallanna et al., 2016) and *CLDN18* has been confirmed as a GATA6 target in gastric cancer (Sulahian et al. 2014), therefore we used these likely targets to test the regulatory potential of HNF4A and GATA6 in OE19 cells. Both of these genes exhibit multiple regions of increased chromatin accessibility in both tumour tissue and OE19 cells, several of which contain motifs for HNF4A or GATA6 binding (Supplementary Fig. S4D). ChIP-qPCR confirmed occupancy of regions by HNF4A and GATA6 containing their cognate binding motifs, while control regions lacking these motifs showed little evidence of binding (Supplementary Fig. S4E).

Next, to gain a more comprehensive view of the role of HNF4A and GATA6 we expanded our analysis of their cistromes by ChIP-seq in OE19 cells. The HNF4A and GATA6 antibodies robustly precipitated the respective proteins (Supplementary Fig. S5A) and replicate ChIP-seq experiments were highly reproducible (Supplementary Fig. S5B and C). We therefore took the overlap of the two replicates forward for further analyses, resulting in 6,870 and 37,658 binding regions for HNF4A and GATA6 respectively (Supplementary Table S3). To ensure that all peaks taken forward were functional, we ranked peaks from both datasets by enrichment, partitioned peaks into 10% bins and searched for respective motifs. All bins showed a high enrichment of motifs ranging from 87% to 32% for HNF4A and 70% to 28% for GATA6 (Supplementary Fig. S5D), therefore we kept all peaks for further analyses. Very similar genomic distributions were observed for each transcription factor, with the majority (90%) being in intra-or intergenic regions (Fig. 2A). DNA motif analysis identified HNF4A and GATA6 as the top enriched motifs in their respective datasets (>42% of regions in both cases)(Fig. 2B) and although these motifs tended to be more prevalent in the higher confidence binding regions, these motifs were distributed throughout the entire datasets (Supplementary Fig. S5D). Interestingly, we identified AP-1 as the next most enriched motif in both cases, in keeping with our previous finding of AP1 as an important factor in OAC (Britton et al., 2017), but in addition we also identified GATA motifs in the HNF4A binding regions and Forkhead binding motifs in the GATA6 binding regions (Fig. 2B; Supplementary Table S4). Although the HNF4A motif was not among the top enriched motifs in the GATA6 ChIP-seq dataset, searching for the HNF4A DNA binding motif within GATA6 bound regions revealed that 8.1% of GATA6 ChIP-seq regions contain a HNF4A motif (Supplementary Fig. S5E). These observations are in keeping with our identification of the same motifs in the context of open chromatin regions in OAC (Fig. 1F) and suggest an integrated network of transcription factors. Indeed, the majority (74%) of HNF4A binding regions are also occupied by GATA6 (Fig. 2C and D). One locus showing such co-occupancy is *IRAK2*, although uniquely bound regions can be identified in other loci including *CLDN18* and *HES4* (Fig. 2E). The regions co-occupied by both HNF4A and GATA6 exhibit low levels of open chromatin in non-cancerous oesophageal Het1A cells, but show elevated levels of open chromatin in OE19 cells compared to the single occupied regions, suggesting that these may have more regulatory potential (Fig. 2D).

**Figure 2.**
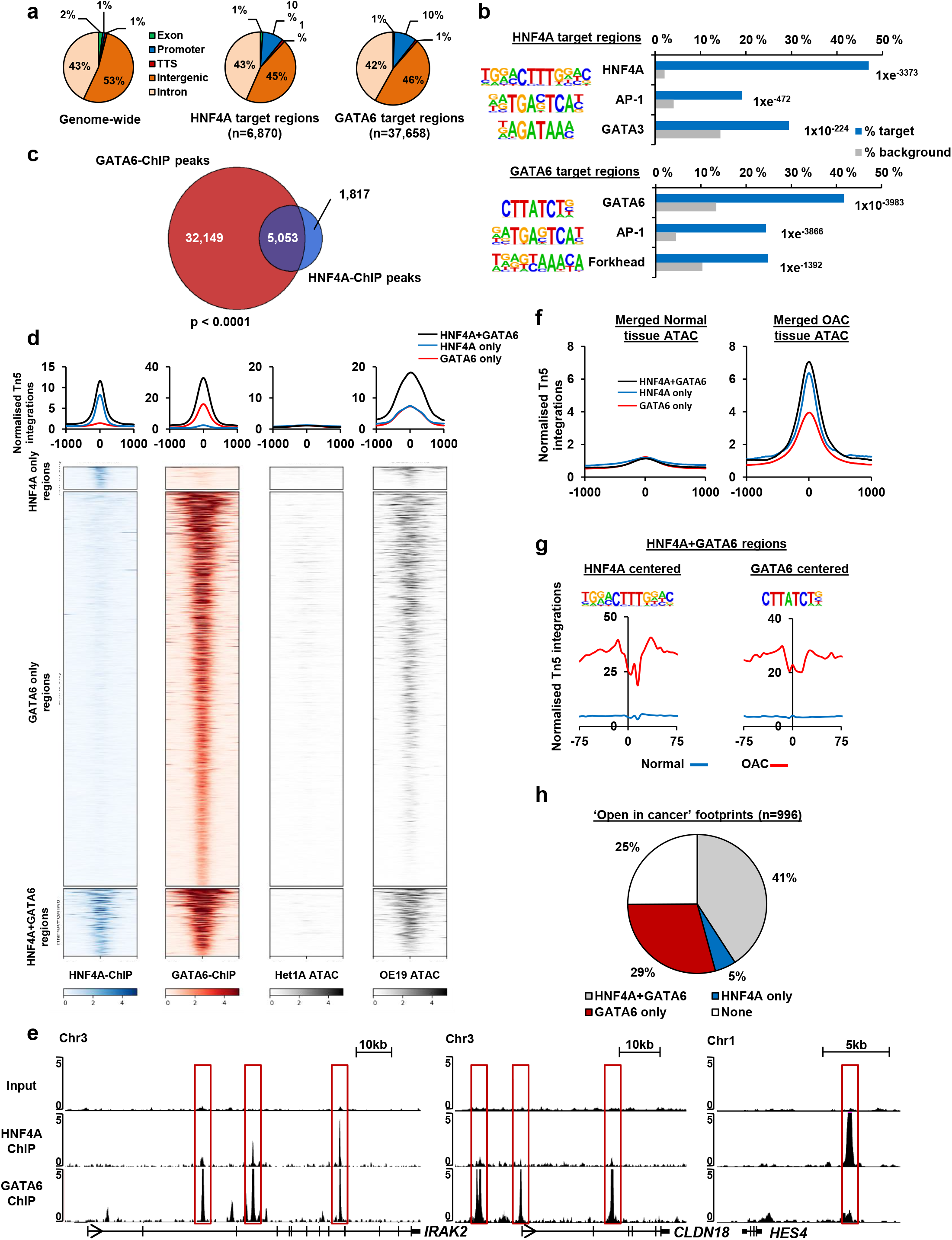
Genome-wide identification of HNF4A and GATA6 binding sites. (A) Genomic distribution of HNF4A and GATA6 ChIP-seq regions compared to genomic average. Promoter is defined as -2.5 kb to +0.5 kb relative to the TSS. (B) Top three motifs from de novo motif discovery and their associated transcription factor found in ChIP-seq regions for HNF4A (left) and GATA6 (right). The frequency of motif occurrence is shown and motifs are sorted by p-value. (C) Venn diagram of overlapping binding regions from HNF4A and GATA6 ChIP-seq. P-value was calculated using Fisher’s exact test. (D) Heatmap and tag density plots of ChIP-seq signals for HNF4A and GATA6 at regions bound by HNF4A only, GATA6 only or both HNF4A and GATA6. The ATAC-seq signals at the same regions in Het1A and OE19 cells are shown. (E) UCSC browser tracks showing ChIP-seq tracks at three loci (*IRAK2, CLDN18* and *HES4*). GATA6 and/or HNF4A bound regions are boxed. (F) Tag density plots of ATAC-seq signal at regions bound by HNF4A only, GATA6 only or both HNF4A and GATA6 in normal and OAC tumour tissue. (G) Footprinting of ATAC-seq data from normal and OAC tissue (bottom) on 150 bp regions surrounding the HNF4A or GATA6 transcription factor motifs found in HNF4A+GATA6 co-occupied ChIP-seq regions. (H) Distribution of ATAC-seq footprints in ‘open in cancer’ regions that coincide with ChIP-seq regions of HNF4A and GATA6.

To relate these findings back to patient-derived samples, we asked whether the HNF4A and GATA6 binding regions are accessible in our biopsies from normal tissue and OAC. Little evidence of open chromatin is apparent in normal tissue but elevated levels are seen in OAC tissue, which is particularly marked in regions occupied by HNF4A alone and co-occupied by both HNF4A and GATA6 in OE19 cells (Fig. 2F). We also asked whether we could detect changes in DNA accessibility in and around the HNF4A and GATA6 binding motifs within their binding regions, when comparing data from normal and OAC samples. Clear footprints were observed centred on the HNF4A motifs in OAC tissue which were particularly prominent in regions co-occupied with GATA6 (Fig. 2G; Supplementary Fig. S5F). Similarly, we also observed a footprint around the GATA6 motifs in OAC tissue but this was only observed in the co-occupied regions (Fig. 2G; Supplementary Fig. S5F). Similar results were obtained when comparing chromatin accessibility data from cell line models (Supplementary Fig. S5F). Finally, we asked whether any of the footprints we identified in the patient-derived OAC material overlapped with the binding regions for HNF4A and GATA6 and found that 75% of these overlapped with one or other factor, with the largest percentage (41%) being associated with the regions co-bound by HNF4A and GATA6 (Fig. 2H).

Collectively, these data demonstrate that HNF4A and GATA6 bind to a large number of regulatory regions in OAC cells. More importantly, these factors usually co-occupy the same regions and these co-occupied sites are potentially more functionally relevant in OAC as they are associated with more open chromatin and deeper footprints in both cell line models and patient samples.

### The GATA6 and HNF4A regulome

Having established the HNF4A and GATA6 cistromes, we next determined their effects on gene expression by depleting them individually and in combination in OE19 cells. Efficient depletion of both transcription factors was achieved at both the RNA and protein levels (Supplementary Fig. S6A-D), and the expected down regulation of the target genes *CLRN3* and *CLDN18* was achieved by depletion of HNF4A and GATA6 respectively (Supplementary Fig. S6B and D). RNAseq was then performed, data quality verified (Supplementary Fig. S6E) and differentially expressed genes identified. We focussed on likely direct target genes (defined as being associated with an annotated ChIP-seq peak) and identified 489 and 1,122 genes whose expression was changed by >1.3 fold (p-value <0.05) following HNF4A and GATA6 depletion respectively (Fig. 3A; Supplementary Table S5A and B). As expected for activating transcription factors, the majority (~70% in both cases) were downregulated. Surprisingly, given the large overlap in binding the overlap in genes down or up regulated by each protein was relatively modest (albeit very significant), suggesting that either HNF4A or GATA6 might be the more dominant factor at different genes (Fig. 3B). To identify further potential regulated genes we performed RNAseq following depletion of both HNF4A and GATA6. This identified a further 156 deregulated genes (69 down and 87 up regulated) when both factors are depleted together in addition to the 92 genes which are commonly reregulated by either treatment alone (Fig. 3C; Supplementary Table S5C). GO term analysis of all the genes downregulated by depletion of either HNF4A or GATA6 showed enriched GO terms for “Glycoprotein metabolic process”, “Response to wounding” and “Regulation of hormone levels” (Fig. 3D). Similar terms were uncovered when individually analysing the genes deregulated by depletion of either factor alone (Supplementary Fig. S6F). Importantly, several of these GO terms are also similar to the GO terms identified in regions that are specifically accessible in patient-derived OAC samples (Supplementary Fig. S1C) indicating the HNF4A and GATA6 regulome contributes to the overall phenotype of OAC.

**Figure 3.**
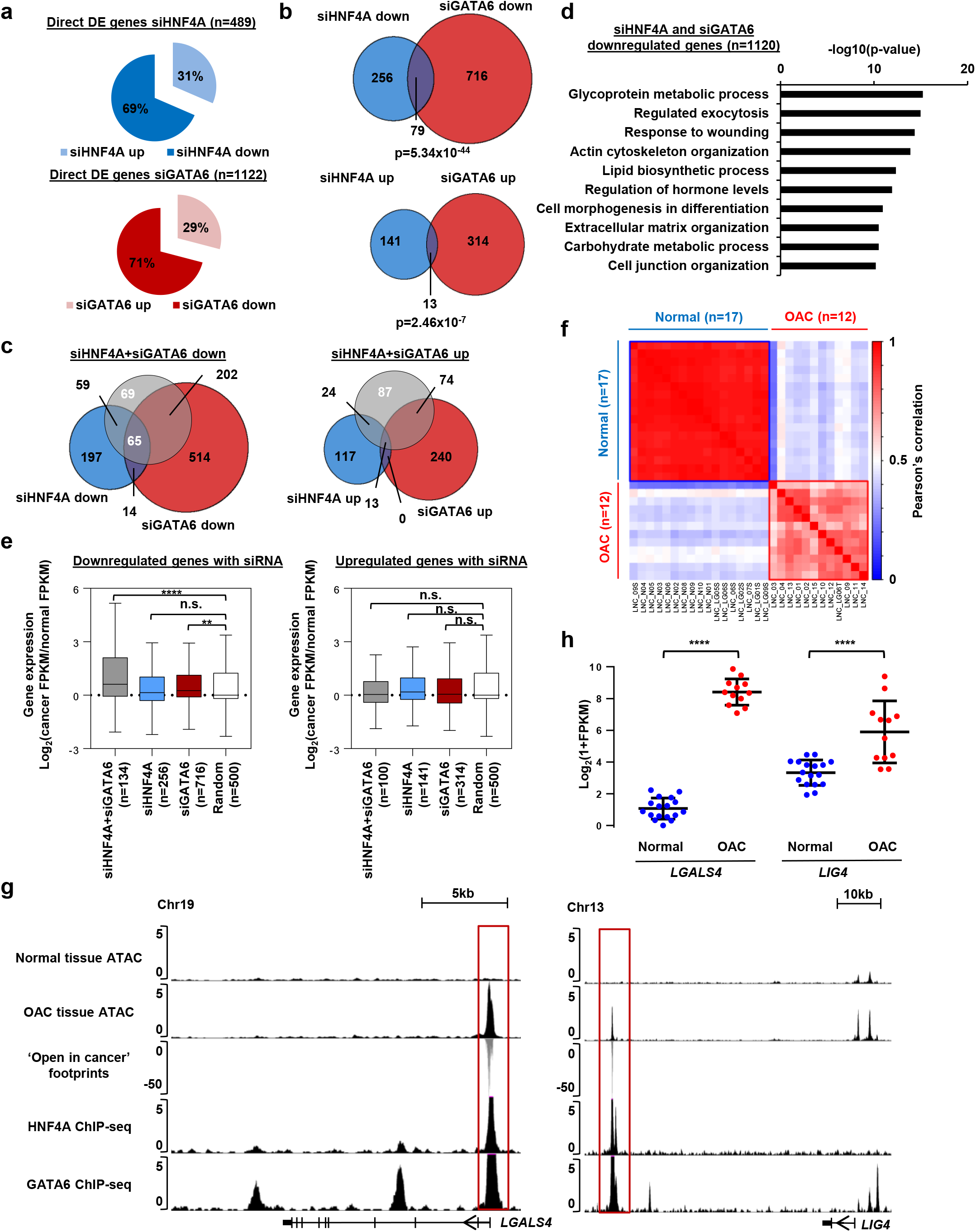
HNF4A and GATA6 target genes are over-expressed in OAC tumours. (A) Proportion of differentially expressed (DE) genes (>1.3 fold; p-value<0.05) that are up and down regulated in OE19 cells following siHNF4A (top) and siGATA6 (bottom) treatment. (B) Overlap of differentially expressed genes in siHNFA and siGATA6 datasets. P-value calculated using the hypergeometric test. (C) Overlap of differentially expressed genes that are downregulated following siHNF4A and siGATA6 co-treatment compared to single siRNA treatments. (D) Top 10 significant “biological processes” GO term analysis of genes downregulated following siHNF4A, siGATA6 or siHNF4A+siGATA6 treatment. (E) Boxplot of changes in gene expression in OAC tissue (shown as cancer/normal FPKM log_2_ fold change) for genes downregulated and upregulated by siRNA against HNF4A (blue), GATA6 (red) or both (grey). Whiskers represent 1.5 x IQR. A random (500 randomly selected RefSeq transcripts) dataset is represented in white. **** represents p<0.0001, ** represents p<0.01. (F) Pearson correlation plot of FPKM values of siHNF4A+siGATA6 regulated genes in normal (n=17) and OAC (n=12) patient samples (Maag et al., 2017). Samples were clustered hierarchically and the two main clusters are highlighted as a ‘normal’ cluster (blue) and a ‘OAC’ cluster (red). (G) UCSC genome browser tracks showing ATAC-seq data in normal and tumour tissue (top), ‘open in cancer’ footprints (centre) and HNF4A and GATA6 ChIP-seq signals (bottom) at genes regulated by HNF4A and GATA6, *LGALS4* and *LIG4*. (H) Plot showing the expression of *LGALS4* and *LIG4* in normal (blue) and OAC (red) tissue samples (Maag et al., 2017). Mean is represented by black bar with standard deviation shown above and below. **** represents p<0.0001.

To determine whether the HNF4A- and GATA6-regulated genes are relevant to OAC, we examined whether any changes in their expression could be observed in OAC compared to normal oesophageal tissue. We focussed on the genes that were downregulated following siRNA treatment as these are more likely direct targets normally activated by these transcription factors. Significantly higher expression of the genes activated by both HNF4A and GATA6 in OE19 cells was observed in OAC tissue, with a lower but still significant increase in expression for the cohort of genes activated by GATA6 alone (Fig. 3E). In contrast, the genes that are upregulated following siRNA treatment (i.e. likely indirect effects) are not expressed at higher levels in OAC tissue (Fig. 3E). An identical trend was observed in a different dataset, with highest expression in cancer being observed for genes activated by both HNF4A and GATA6 (Supplementary Fig. S7). Hierarchical clustering using Pearson’s correlation of the expression of genes activated by both factors in normal and OAC tissue completely separates the two tissues (Fig. 3F), again suggesting that the expression of HNF4A, GATA6 and their target genes are biologically relevant. Two example genes from this category are *LGALS4* and *LIG4* (Fig. 3G). Both are direct targets for HNF4A and GATA6, both are associated with regions of open chromatin around these sites only in OAC tissue, and both have OAC-specific footprints. Importantly, these changes in HNF4A and GATA6 binding activity are also associated with increased gene expression in the context of OAC (Fig. 3H).

Collectively, these data demonstrate that HNF4A and GATA6 directly regulate a set of genes that are expressed at higher levels in OAC tissue, and are consistent with our identification of HNF4A and GATA6 binding motifs in the open regulatory regions that are specific to OAC.

### The HNF4A-GATA6 regulatory network is operational in Barrett’s oesophagus

Our results suggest an important role for the regulatory network involving HNF4A and GATA6 in OAC. However, OAC is thought to usually arise from a pre-cancerous metaplastic state known as Barrett’s oesophagus (Desai et al., 2012). We therefore asked whether this regulatory network could be detected in Barrett’s oesophageal tissue. First we performed ATAC-seq on non-dysplastic Barrett’s tissue taken from four different patients. The data from these samples was highly consistent (Supplementary Fig. S8A). PCA analysis demonstrated that the Barrett’s samples clustered together with the OAC samples rather than the samples from normal tissue (Fig. 4A). We therefore compared the open chromatin regions from Barrett’s samples with those found in normal oesophageal tissue (Supplementary Table S6). The majority (>98%) of changes in accessibility were found in non-promoter regions i.e. intra- and intergenic regions with chromatin opening being the predominant (64%) change in Barrett’s cells (Fig. 4B and 4C). Interestingly, “gland development” was among the enriched GO terms in genes associated with opening chromatin regions, which is in keeping with the conversion of the stratified epithelium of the oesophagus to a glandular epithelium in Barrett’s cases (Supplementary Fig. S8C).

**Figure 4.**
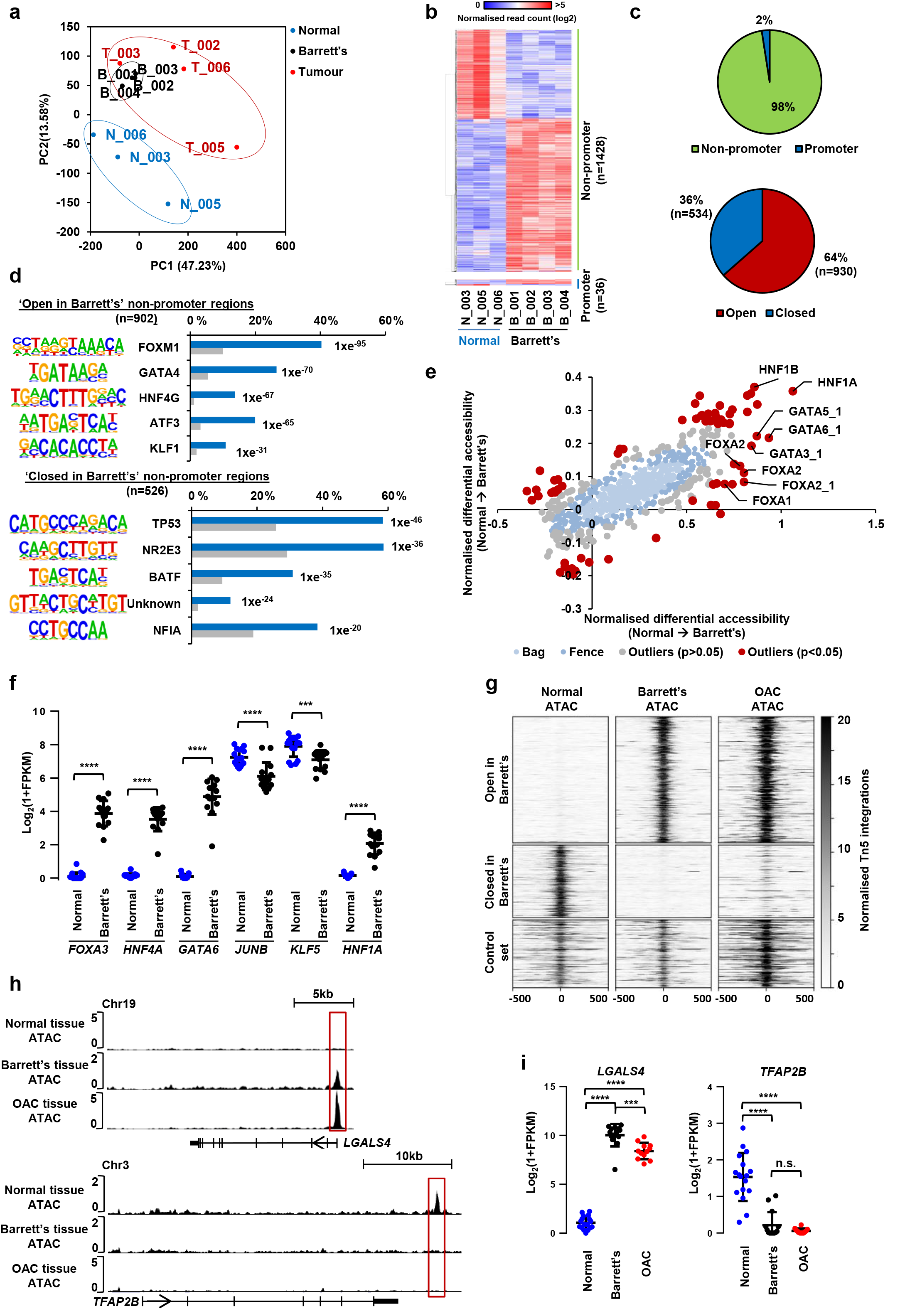
Barrett’s oesophagus ATAC-seq is related to OAC. (A) PCA plot of the normalised ATAC-seq signal across the top 50,000 regions of all 3 normal (blue), 4 Barrett’s (black) and 4 OAC (red) tissue samples. (B) Heatmap of normalised Tn5 cleavage events in a +/− 250 bp region surrounding the summit of differentially accessible non-promoter and promoter regions (>5 fold change, p<0.05) in normal vs Barrett’s samples. (C) Pie charts representing the proportion of differentially accessible regions that are located in promoter and non-promoter regions (top) and regions that are five-fold more open or closed in Barrett’s tissue (bottom) (D) Top five DNA motifs derived from *de novo* motif discovery and their associated transcription factor that are enriched in ‘open in Barrett’s’ (top) or ‘closed in Barrett’s’ (bottom) non-promoter regions. The frequency of motif occurrence is shown and the motifs are sorted by p-value. (E) Scatter bag plot of differential chromatin accessibility (x-axis) and footprinting (y-axis) depth around human transcription factor binding motifs in normal and Barrett’s tissue. Significant outliers (p<0.05) are represented in red. Motifs with enrichment in “Open in Barrett’s” and “Closed in Barrett’s” regions are labelled. (F) Plot of Log_2_ FPKM values of transcription factors with enriched motifs in differentially accessible regions from normal (blue) vs Barrett’s analysis (“open in Barrett’s”) in RNAseq data from normal or Barrett’s (black). Mean is represented by black bar with standard deviation shown above and below. **** represents p<0.0001, *** represents p<0.001. (G) Heatmap of ATAC-seq signal from normal, Barrett’s and OAC tissue at differentially “Open in Barrett’s”, “Closed in Barrett’s” and a control set of 500 random nonsignificantly differential accessible regions. (H) UCSC genome browsers tracks of ATAC-seq signal at two example loci (*LGALS4* and *TFAP2B*) with differentially accessible regions boxed. (I) Expression of *LGALS4* and *TFAP2B* in normal, Barrett’s and OAC tissue (Maag et al., 2017). Mean is represented by black bar with standard deviation shown above and below. **** represents p<0.0001, *** represents p<0.001.

Next, to identify potential upstream regulatory proteins, we searched for enriched motifs in ‘open in Barrett’s’ non-promoter regions and identified motifs for Forkhead, GATA and HNF4 related transcription factors as the most enriched in the regions which are more accessible in Barrett’s cells. Conversely, the p53 binding motif is enriched in the regions which become less accessible in Barrett’s cells (Fig. 4D; Supplementary Table S7). We also used BaGFoot (Baek et al., 2017) to identify motifs that showed evidence of changes of occupancy through altered “footprinting” in all of the open chromatin regions in the Barrett’s samples. Again, we identified motifs for GATA6 and Forkhead transcription factors and in addition the motifs for HNF1A/B showed particularly strong increases in both footprinting depth across the motif and accessibility in the local surrounding area (Fig. 4E; Supplementary Fig. S9). Although the HNF4A motif was not identified using BaGFoot, centring accessible regions in Barrett’s onto the HNF4A motif and plotting ATAC-seq signal across these regions shows a clear footprint in Barrett’s cells compared to normal tissue (Supplementary Fig. S8D). It therefore appears that the same transcription factors which we identified in OAC are associated with the open chromatin regions in Barrett’s oesophageal cells. To identify the likely transcription factors binding to these sites, we examined the expression of several family members in Barrett’s and normal oesophageal tissue (Maag et al., 2017). GATA6, HNF4A/G, HNF1A/B and FOXA1,2,3 are all expressed to higher levels in Barrett’s samples (Fig. 4E; Supplementary Fig. S8D). Given the prominent appearance of HNF4A and GATA6 binding motifs, we examined the expression of the genes in Barrett’s samples that are directly regulated by HNF4A and GATA6 in OE19 cells. In keeping with a likely regulatory role for these transcription factors, these same sets of genes are also upregulated in Barrett’s tissue (Supplementary Fig. S8F).

Finally, as a direct comparison, we compared the open chromatin landscape of OAC cells with that found in Barrett’s and normal oesophageal tissue. Regions that are more accessible in Barrett’s oesophagus cells maintain this accessibility in OAC cells. Closed regions also maintain a similar state of accessibility in OAC (Fig. 4G). Genome browser tracks of the *LGALS4* (gained accessible region) and *TFAP2B* (lost accessible region) loci illustrate the “maintenance” of chromatin accessibility states in OAC compared to Barrett’s oesophagus (Fig. 4H). Expression of *LGALS4* and *TFAP2B* show higher or lower expression in Barrett’s oesophagus respectively, and this level of expression is maintained in OAC, mirroring the loci accessibility patterns (Fig. 4I).

Together this data therefore indicates that the regulatory network involving HNF4A, GATA6, HNF1B and FOXA transcription factors is already established in Barrett’s metaplastic cells and is maintained in OAC cells.

### HNF4A drives the formation of open chromatin

Our results are consistent with two possible models. Either one or more of the transcription factors in the regulatory network can bind to pre-configured open chromatin in Barrett’s and OAC cells or instead they might themselves directly trigger this opening. To test the latter possibility, we used lentivirus with doxycycline inducible constructs to express either HNF4A or GATA6 in the “normal” oesophageal Het1A cells and profiled the resulting open chromatin landscape using ATAC-seq after induction.

Western blot and RT-qPCR analysis demonstrated induction of HNF4A and GATA6 protein and mRNA after 2 days of doxycycline treatment and this was maintained at 4 days (Fig. 5A, Supplementary Fig. S10A). ATAC-seq was then performed after 2 days and 4 days of doxycycline treatment. All replicates had high levels of correlation (Supplementary Fig. S10B), thus we merged the alignment files of the replicates of all samples from either the HNF4A expression or GATA6 expression time course and re-called peaks from each of these combined datasets. We then took the top 50,000 most significant regions to calculate differential accessibility between parental Het1A cells and Het1A-HNF4A and Het1A-GATA6 cells at two and four days of induction (Supplementary Table S8A and S8D). Differential accessibility analysis determined that HNF4A overexpression resulted in 2973 regions with increased accessibility and 976 regions with decreased accessibility after two days of induction. In contrast, after two days of induction of GATA6, only 87 regions became more accessible, and only 149 became less accessible. After four days of induction, the number of differential accessible regions remained similar for both transcription factors, thus we focussed on two days of induction to identify the immediate effects of transcription factor overexpression (Supplementary Fig. S10C; Supplementary Table S8A-F). As HNF4A appears to be able to drive the formation of open chromatin much more robustly that GATA6, we decided to focus on the chromatin regions that are dynamically controlled by HNF4A. The genomic distribution of regions that become more accessible after HNF4A induction is similar to the HNF4A binding regions identified by HNF4A ChIP-seq (Supplementary Fig. S10D), and there was extensive overlap of accessible regions between 2 days and 4 days of induction (Supplementary Fig. S10E). *De novo* transcription factor motif discovery in the chromatin regions that open up following HNF4A expression, uncovered over-representation of binding motifs for AP-1, HNF4A and TEAD (Fig. 5B; Supplementary Table S9). Surprisingly, the GATA motif was not enriched in these regions. We next used BaGFoot to simultaneously look for differential accessibility and footprinting at these regions. The HNF4A motif showed both increased footprinting and increased accessibility (Fig. 5C; Supplementary Fig. S11). These data indicate that the differential accessible regions induced by HNF4A overexpression are indeed driven by HNF4A.

**Figure 5.**
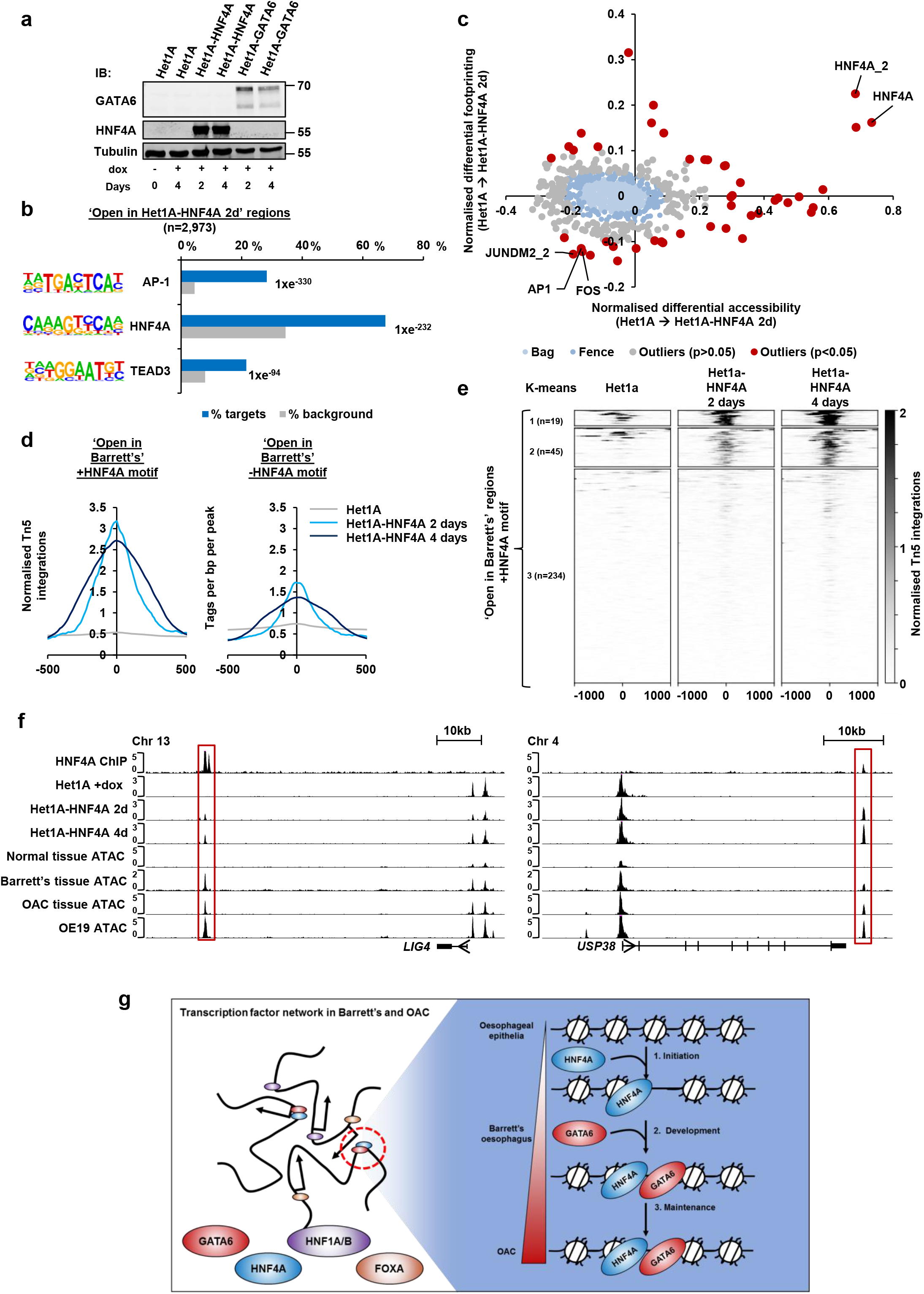
HNF4A demonstrates pioneer factor function. (A) Immunoblot analysis of HNF4A, GATA6 and tubulin in Het1A, Het1A-HNF4A and Het1A-GATA6 cells. The addition of doxycycline for 2 or 4 days is indicated (+). (B) Top three DNA motifs derived from *de novo* motif discovery of differentially accessible regions in Het1A-HNF4A cells after two days of induction and their associated transcription factors. The frequency of motif occurrence is shown and the motifs are sorted by p-value. (C) Scatter bag plot of differential chromatin accessibility (x-axis) and footprinting (y-axis) depth around human transcription factor binding motifs in Het1A and Het1A-HNF4A cells. Significant outliers (p<0.05) are represented in red. Motifs with enrichment in “Open in Het1A-HNF4A”regions are labelled. (D) Tag density plots of ATAC-seq signal from Het1A and Het1A-HNF4A cells (2 days and 4 days doxycycline induction) around differentially open regions in Barrett’s tissue with (left) or without (right) a HNF4A consensus binding motif. (E) Heatmap of ATAC-seq signal from Het1A and Het1A-HNF4A cells (2 days and 4 days doxycycline induction) at “Open in Barrett’s” regions with a HNF4A motif. Regions were subjected to K-means hierarchical clustering (k=3). (F) UCSC genome browser tracks of two example loci at *LIG4* and *USP38* with regions boxed which show increased accessibility with HNF4A overexpression. (G) Model of the transcription factor network in Barrett’s and OAC. HNF4A is able to promote chromatin opening in normal oesophageal cells. In Barrett’s, a transcription factor network, including HNF4A and GATA6 are co-expressed and these two transcription factors co-occupy a large number of genomic regions. This regulome persists in the context of the chromatin landscape of OAC cells.

We next sought out to investigate whether the regions of differential accessibility in Barrett’s are associated with the HNF4A-induced accessible regions in Het1A cells. First we centred the “open in Barrett’s” regions onto their HNF4A motifs and then plotted the average normalised ATAC-seq signal from parental Het1A and Het1A-HNF4A at 2 and 4 days of induction. Regions with a motif showed large increases of accessibility at 2 days and this was maintained at 4 days, whereas regions without a motif showed lower levels of increased accessibility (Fig. 5D). To further delineate the induction of chromatin accessibility, we took all of the regions which showed increased accessibility in Barrett’s, centred them on the HNF4A motif and plotted the tag densities of the ATAC-seq signals from the HNF4A induction profile in Het1A cells around the centre of these peaks. The profiles were then subjected to k-means clustering which revealed three clusters, characterised by strong opening of chromatin (cluster 1), more moderate levels of chromatin opening (cluster 2) and little chromatin opening (cluster 3) (Fig. 5E; Supplementary Fig. S10G). Regions that show evidence of increased accessibility following HNF4A-overexpression make up 64/298 (21%) of the open chromatin regions in Barrett’s with a HNF4A motif. Two example regions are associated with *LIG4* and *USP38*, which show an induction of chromatin opening with HNF4A, a region of accessibility in Barrett’s and OAC and also show direct evidence of HNF4A by ChIP-seq (Fig. 5F). More generally we asked whether the genes associated with HNF4A-mediated open chromatin regions showed any communalities to those found in OAC or Barrett’s tissue samples. Similar biological processes are affected with several GO terms were found to be in common such as “regulation of cell adhesion”, “gland development” and “response to wounding” (Supplementary Fig. S10F).

Collectively, these results indicate that HNF4A is capable of inducing chromatin opening in normal oesophageal cells, and in this context, it is able to promote the opening of regions of chromatin that are seen to be differentially accessible in Barrett’s and OAC. HNF4A therefore has the potential to promote a pivotal initiation event in Barrett’s oesophagus formation.

## Discussion

Cancer genome sequencing efforts have given us insights into the mutational spectrum and hence the potential molecular causes of OAC (Weaver et al., 2014; Dulak et al., 2013; Ross-Innes et al., 2015). These have contributed to a model where OAC develops from a pre-cancerous state known as Barrett’s oesophagus. Barrett’s oesophagus itself is thought to be caused by acid reflux around the gastro-oesophageal junction (GOJ) which triggers a change in the structure of the epithelial lining of the oesophagus from a stratified epithelium to a glandular epithelium (reviewed in Spechler and Souza, 2014). However, in addition to genetic changes, epigenetic alterations to the chromatin landscape are also likely to play an important role in disease progression. Several recent studies that have used ATAC-seq and open chromatin profiling to uncover the regulatory networks involved in cancer such as in small cell lung cancer metastasis (Denny et al., 2016), ER-dependent breast cancer (Toska et al., 2017) and EMT (Pastushenko et al., 2018). Here, we have used ATAC-seq to uncover the regulatory open chromatin landscape of OAC and non-dysplastic Barrett’s samples from patients, and demonstrate that they exhibit many similarities. This provides strong molecular support for Barrett’s being a precursor state to OAC.

Previously we identified AP1 as an important regulatory transcription factor in OAC (Britton et al., 2017). Here we took a more focussed approach and investigated a subset of OAC samples which enabled us to uncover a regulatory network comprised of HNF4A, GATA6, HNF1B and FOXA1 transcription factors that exists in OAC but is already activated in Barrett’s (Fig. 4D and 4E; Fig. 5G). These transcription factors have all been shown to play a role in intestinal development (Verzi et al., 2010; Beuling et al., 2012; Kerschner et al., 2014; Walker et al., 2014; Gosalia et al., 2015; San Roman et al., 2015; Yang et al., 2016), suggesting that Barrett’s and OAC cells have reverted to a more primitive state where these factors are operational. For example, HNF4A, GATA6 and FOXA3 are all broadly expressed in the stomach, liver, pancreas and intestine during mouse embryogenesis but are absent from the oesophagus (Sherwood et al., 2009). Similarly, we show that these factors are all expressed during early human intestinal development but with the exception of early definitive endoderm, co-expression is not observed in any particular organ (see Supplementary Fig. S3). An alternative hypothesis is that rare cells may exist that contain this regulatory network and may act for example as stem cells for replenishing the oesophageal epithelium. By focussing on two of the highest expressed members of this network, HNF4A and GATA6 we showed that these two factors co-occupy many genomic loci and work together to control gene expression. However, it is likely that a much more complicated interacting network exists with multiple combinatorial interactions involving these two transcription factors, HNF1B and FOXA1. Indeed, in addition to the family members we have focussed on, other related proteins such as HNF4G, GATA4, HNF1A, and FOXA2/3 are also expressed at higher levels in Barrett’s and OAC. Thus it is likely that there is functional redundancy built into the regulatory network, which may in part explain why the numbers of genes we find to be regulated by HNF4A and GATA6 is an order of magnitude lower than the number of direct targets. It is currently unclear how this network is initially established but we demonstrate that HNF4A is able to penetrate and open up regions of inaccessible chromatin which may represent one of the initiating events for Barrett’s development. Indeed, these results are consistent with the observation that HNF4A over-expression in mouse oesophageal epithelial cells is sufficient to induce the expression of several Barrett’s-specific markers (Colleypriest et al., 2017). This activity of HNF4A is akin to pioneering activity which has been demonstrated for several transcription factors (Zaret and Carroll, 2011). However, GATA6 did not demonstrate widespread pioneering activity, and there is a distinct lack of GATA motifs in the HNF4A-induced accessible chromatin regions, despite our demonstration of widespread co-binding of HNF4 and GATA factors in Barrett’s and OAC cells. This suggests that although HNF4A may be an initial driver, additional events must occur to facilitate subsequent GATA factor binding to expand the regulatory network.

GATA4 and GATA6 have previously been implicated in OAC by the numerous studies which have observed that their loci are frequently amplified in the transition from Barrett’s oesophagus to cancer (Lin et al., 2012; Stachler et al., 2015). However, we show here that these transcription factors are already expressed in Barrett’s and are associated with largely the same open chromatin regions in both Barrett’s and OAC. This suggests that the supra physiological levels of GATA4/6 arising from genomic amplifications, may act on a different set of loci to drive OAC formation. Patient samples containing such amplifications are needed to test this hypothesis. Interestingly, GATA6 has also been shown to operate in gastric cancer (Sulahian et al., 2014; Chia et al., 2015). Recent genome sequencing data indicated that CIN variant gastric, GOJ and OAC tumours are closely related at the molecular level (The Cancer Genome Atlas Network, 2017). Our data therefore further support this conclusion. Indeed, the OE19 cells we use here to functionally validate a role for HNF4A and GATA6 were isolated from the GOJ, and these cells contain an open chromatin landscape that most closely resembles the data from patient–derived OAC samples.

HNF1A is known to respond to bile acids and has been shown to upregulate *MUC4* in OAC cells (Piessen et al., 2007). Furthermore, HNF4A has previously been implicated in the initial response of normal human oesophageal mucosa to bile acids (Green et al., 2014). More recently, HNF4A has been shown to be sufficient to induce a columnar-like phenotype in adult mouse oesophageal epithelium (Colleypriest et al., 2017). These results suggest that HNF1A/B and HNF4A may play an initiating role in promoting the transition to Barrett’s in human disease. Indeed, our results demonstrate that forced expression of HNF4A in normal oesophageal epithelial cells is sufficient to trigger chromatin opening that are observed in Barrett’s and maintained in OAC. This provides further evidence to suggest that Barrett’s might arise directly from the oesophageal squamous epithelium. Alternatively, the same mechanisms might trigger the transition from other cell types that have been proposed as the cell of origin including cells from the GOJ (Jiang et al., 2017) or migrating cells of gastric origin (Quante et al., 2012).

While we have uncovered a transcriptional regulatory network that clearly links OAC cases to underlying Barrett’s, it is possible that cells may directly transition to OAC without transitioning through Barrett’s. Indeed two of the OAC cases that we have studied do not possess the open chromatin landscape that is characteristics of Barrett’s oesophageal cells (Supplementary Fig. S1A; Britton et al., 2017). Further samples are needed to study alternative open chromatin landscapes and the underlying regulatory networks that might be established in OAC

In summary, we have used open chromatin profiling to uncover the transcriptional regulatory networks that are operational in Barrett’s oesophagus and retained in OAC. This provides molecular insights into the stepwise progression towards OAC and implicates the re-activation of a set of transcription factors usually associated with primitive intestinal development from the HNF4, GATA, FOXA and HNF1 subfamilies. This is therefore a potentially powerful approach to uncover regulatory pathways in cancer cells, stratify cancers, and identify biomarkers which can complement and extend the insights being provided through ongoing genomic sequencing efforts.

### Experimental procedures

#### Ethics statement

Ethical approval for collection of Barrett’s oesophagus tissue samples from patients at Leigh Infirmary was granted by the ethics committees Salford Royal NHS Foundation Trust (2010) respectively (04/Q1410/57). Patient consent was obtained in written form and signed by the patient and doctor.

### Cell culture

All cells were cultured in an incubator maintained at 37°C and 5% CO_2_. OE19 cells were cultured in RPMI 1640 media (Gibco) supplemented with 10% FBS (Gibco) and 1% penicillin-streptomycin (Gibco). Het1A and HEK293T cells were cultured in DMEM media (Gibco) with 10% FBS and 1% penicillin-streptomycin. Human tissue was processed as described previously (Britton et al., 2017). The expression of HNF4A and GATA6 was induced by treating Het1A-HNF4A or Het1A-GATA6 cells with 100 ng/mL doxycycline (Sigma). Parental Het1A cells were treated with 100ng/mL doxycycline as a control for 4 days.

### Barrett’s oesophagus tissue collection

~2mm Barrett’s oesophagus samples were obtained from four consenting patients. All patients presented with at least C3M3 Barrett’s, which is defined as ‘long’ Barrett’s and is of higher risk of progression to OAC (Fitzgerald et al., 2014). Intestinal metaplasia histology and no evidence of dysplasia was confirmed for all samples.

### RNA isolation and RT-qPCR

RNA was isolated from 2 × 10^5^ cells using a RNeasy RNA extraction kit (Qiagen, 74136) as per the manufacturer’s instructions. RT-qPCR was carried out using QuantiTect SYBR^®^ Green RT-PCR Kit (Qiagen) using the primer pairs detailed in Table S10. Relative gene expression was calculated using the ΔΔCT method relative to levels of *RPLP0* mRNA.

### siRNA transfection

SMART-pool siRNAs against *HNF4A* (Dharmacon, L-003406-00-0005), *GATA6* (Dharmacon, L-008351-00-0005), and control non-targting siRNA (siNT; Dharmacon, D-001810-10-0020) were transfected using Lipofectamine^®^ RNAiMAX Transfection Reagent (Thermofisher, 13778150) as per the manufacturer’s instructions. Briefly, 2 × 10^5^ cells were transfected with 100 pmol siRNA and RNA was harvested 72 hours later.

### RNA-seq analysis

RNA-seq libraries were prepared using a TruSeq stranded mRNA sample prep kit and run on a HiSeq 4000 (Illumina) platform. Illumina adapters were trimmed using Trimmomatic v0.32 (Bolger et al., 2014). Trimmed reads were aligned to the RefSeq transcript annotation of human genome 19 (hg19) using STAR aligner v2.3.0e (Dobin et al., 2013). Reads mapping to chromosomes 1-23, X and Y were retained. Gene expression levels (FPKMs) were estimated using CuffNorm from the Cufflinks package v2.2.1 (Trapnell et al., 2012). Differential expression analysis was carried out using CuffDiff from the Cufflinks package v2.2.1 with default settings.

### ChIP and ChIP-seq analysis

ChIP and ChIP-seq was carried out as described previously (Wiseman et al., 2015). For ChIP-qPCR, 5 × 10^6^ cells, 20 μl appropriate Dynabeads™ (Protein A beads for rabbit IgG and Protein G beads for mouse IgG; ThermoFisher, 10004D/10002D) and either 1 μg HNF4A antibody (R&D Systems, PP-H1415-00), GATA6 antibody (CST, D61E4 XP) was used. PCR was carried out using the primer pairs detailed in Table S10. For ChIP-seq, 1 × 10^7^ cells, 2.5 μg antibody and 50 μl Dynabeads™ were used. Either normal rabbit IgG (Millipore, 12-370) or normal mouse IgG (Millipore, 12-371) antibody was used in parallel as control experiments. DNA libraries were prepared using TruSeq ChIP sample prep kit (Illumina) and sequenced on a HiSeq 4000 (Illumina) platform. Sequencing reads were aligned to NCBI Build hg19 using Bowtie2 v2.2.3 (Langmead et al., 2009). Only reads with a mapping quality >q30 were retained. Peak calling was performed on individual replicates using MACS2 v2.1.1 (Zhang et al., 2008) using default parameters with additional – SPMR parameter. The overlap of peaks between two independent biological datasets was calculated using BEDtools v2.26.0 (Quinlan and Hall, 2010) using bedtools intersect with default settings and using replicate one as a reference. Peaks present in both datasets were taken forward for further analysis.

### ATAC-seq and analysis

ATAC-seq was performed as described previously (Britton et al., 2017). Differential accessibility was determined by merging alignment files of all datasets from both conditions and peaks recalled using MACS2 v2.1.1 (Zhang et al., 2008). The top 50,000 regions ranked by q-value were taken forward, a region +/− 250bp was taken around the peak summit and regions were separated by promoter (−2.5kb,+0.5kb) and non-promoter regions. Promoter and non-promoter regions were tested for differential accessibility using CuffDiff from the Cufflinks package v2.1.1 (Trapnell et al., 2012). Differential accessible regions with a linear fold change of >5 and a p-value of <0.05 were considered significant and were subject to further analyses.

### Bioinformatics analysis

To visualise ATAC-seq and ChIP-seq tag densities, normalised tags were counted using HOMER v4.7 annotatePeaks.pl with –hist parameter (Heinz et al., 2010) and plotted in Microsoft Excel. Chromatin accessibility and gene expression heatmaps at individual loci/genes were drawn using Morpheus (https://software.broadinstitute.org/morpheus/) and hierarchical clustering was performed with this software using 1-Pearson’s correlation unless otherwise stated. ChIP-seq heatmaps and correlation plots were drawn using DeepTools v2.5.2 (Ramírez et al., 2016). *De novo* motif analysis was carried out using HOMER v4.7 findMotifsGenome.pl with –cpg and –mask parameters and motif counting within regions was also carried out using HOMER v4.7 annotatePeaks.pl with –m parameter (Heinz et al., 2010). ATAC-seq and ChIP-seq gene annotation to hg19 was performed using HOMER annotatePeaks.pl, implementing the closest gene model. Gene ontology analysis was carried out using Metascape (metascape.org; Tripathi et al., 2015). Principal component analysis scores were calculated using prcomp in R from ATAC counts in the top 50,000 accessible regions ranked by q-value and were plotted using Microsoft Excel. Correlation plots were generated using R package Corrplot (Wei and Simko, 2017; https://github.com/taivun/corrplot). Footprinting analysis on a subset of ATAC peaks was carried out using the Wellington footprinting algorithm in the pyDNase package v0.2.4 (Piper et al., 2013) and genome-wide simultaneous differential accessibility and footprinting analysis was carried out using BaGFootR v0.9.7 (Baek et al., 2017).

### Protein isolation and analysis

Protein was isolated from 2 × 10^5^ cells by lysing cells in RIPA buffer on ice and sonicating for 5 minutes 30s on/off (Diagenode bioruptor). Protein was quantified using Pierce^™^ BCA Protein Assay Kit (ThermoFisher, 23227). Equal amounts were run on 10% poly-acrylamide gel electrophoresis gels and transferred to Amersham Protran nitrocellulose membrane (GE life sciences, 1060002). Membranes were blocked in Odyssey^^®^^ Blocking Buffer (PBS) (Licor, 927-4000), probed with anti-HNF4A (R&D Systems, PP-H1415-00), anti-GATA6 (CST, D61E4 XP), anti-Tubulin (Sigma, T9026) antibodies with IRDye^^®^^ secondary antibodies (Licor, 925-32212, 925-32213) and scanned with Odyssey^^®^^ IR scanner (Licor).

### Lentivirus vectors, transduction and stable cell line production

A lentiviral vector containing HNF4A was generated by amplifying HNF4A from pcDNA3.1-HNF4A8 (a gift from Frances Sladek; Vuong et al., 2015) with primers containing restriction enzyme sites for BamHI and EcoRI (5’-GATCGGATCCCCCACCATGGTCAGCGTG-3’, ADS6352;5’-GATCGAATTCCGCTAGATAACTTCCTGCTT-3’, ADS6353). The PCR fragment was inserted into pENTR1A and transferred to pInducer20 by Gateway cloning to create pAS4393. A lentiviral vector containing GATA6 (pSLIKneo 3XFLAG-wtGATA6-3XAU1_RNAi resistant) was a gift from Kevin Janes (Addgene, 72616; Chitforoushzadeh et al., 2016). The BleoR gene was amplified from this plasmid using primers containing restriction enzyme sites for XhoI and PacI (5’-GATCCTCGAGGTGTGTCAGTTAGGGTGTGG-3’, ADS6354; 5’-GATCTTAATTAATCGAAATCTCGTAGCACGTG-3’, ADS6355) and inserted back into pSLIKneo 3XFLAG-wtGATA6-3XAU1_RNAi resistant using XhoI and PacI sites to create pAS4395. Lentiviruses were produced as previously described (Nowicki-Osuch et al., 2017). Briefly, HEK293T cells were transfected with pMD2.G (Addgene, 12259), psPAX2 (Addgene, 12260) and the target plasmid using Polyfect (Qiagen, 301107). Viral particles were precipitated from media using PEG-it (System Biosciences, LV810A-1) and Het1A cells were transduced by viral particles using Polybrene (EMD Millipore, TR-1003) as the transduction reagent. Het1A cells containing viral particles were selected with either G418 for Het1A-HNF4A cells (500 μg/mL) or Zeocin for Het1A-GATA6 cells (300 μg/mL) for 14 days to ensure that all non-transduced cells were dead. Selected cells were maintained in DMEM containing either 250 μg/mL G418 (ThermoFisher, 10131027) or 100 μg/mL Zeocin^™^ (ThermoFisher, R25001).

### Statistical analysis

To determine statistical significance between two groups, a Student’s unpaired two-tail T-test was carried out using GraphPad Prism v7. To assess the significance of motif co-occurance distributions, a Chi-squared test was carried out in GraphPad Prism v7. To assess the significance of gene/region overlaps, a hypergeometric distribution test was carried out using the phyper function in R. P-values <0.05 were considered as significant.

### Data access

Data generated in this study are available at ArrayExpress (ATAC-seq data from Barrett’s tissue, E-MTAB-6751; HNF4A and GATA6 ChIP-seq data from OE19 cells, E-MTAB-6858; RNAseq data for HNF4A and/or GATA6 knockdown in OE19 cells, E-MTAB-6756; ATAC-seq data from Het1A (+dox), Het1A-HNF4A and Het1A-GATA6 (2d and 4d), E-MTAB-6931). Human oesophagus, Barrett’s, high-grade dysplastic Barrett’s and OAC RNA-seq was sourced from ArrayExpress (E-MTAB-4054)(Maag et al., 2017). Additional OAC RNA-seq data was obtained directly from TCGA (https://tcga-data.nci.nih.gov/docs/publications/esca_2017/; Cancer Genome Atlas Research Network, 2017). ATAC-seq data from human oesophageal tissue (normal and OAC), and oesophageal cell lines was obtained from E-MTAB-5169 (Britton et al., 2017) and E-MTAB-4209 (Garcia et al., 2016).

## Acknowledgements

We thank Mairi Challinor for excellent technical assistance; Frances Sladek and Kevin Janes for reagents and staff in the Genomic Technologies and Bioinformatics Core Facilities. We also thank Munazah Andrabi for bioinformatics advice and guidance, and Dave Gerrard for the analysis of human embryonic RNAseq datasets. We also thank James Britton for collection of Barrett’s oesophagus samples. We thank Xiaodun Li and Rebecca Fitzgerald on behalf of the OCCAMS consortium for the normal and tumour sample collection. Thanks to Shen-Hsi Yang, Dave Gerrard and Nicoletta Bobola for critical appraisal of the manuscript. This work was funded by grants from the Wellcome Trust, CRUK (through the MCRC) and an MRC DTP studentship to SW.

## References

Baek, S., Goldstein, I., and Hager, G.L. (2017). Bivariate Genomic Footprinting Detects Changes in Transcription Factor Activity. Cell Rep. 19, 1710–1722.

Bao, X., Rubin, A.J., Qu, K., Zhang, J., Giresi, P.G., Chang, H.Y., and Khavari, P.A. (2015). A novel ATAC-seq approach reveals lineage-specific reinforcement of the open chromatin landscape via cooperation between BAF and p63. Genome Biol. 16, 284.

Beuling, E., Aronson, B.E., Tran, L.M.D., Stapleton, K.A., ter Horst, E.N., Vissers, L.A.T.M., Verzi, M.P., and Krasinski, S.D. (2012). GATA6 Is Required for Proliferation, Migration, Secretory Cell Maturation, and Gene Expression in the Mature Mouse Colon. Mol. Cell. Biol. 32, 3392–3402.

Bolger, A.M., Lohse, M., and Usadel, B. (2014). Trimmomatic: A flexible trimmer for Illumina sequence data. Bioinformatics 30, 2114–2120.

Britton, E., Rogerson, C., Mehta, S., Li, Y., Li, X., Fitzgerald, R.C., Ang, Y.S., and Sharrocks, A.D. (2017). Open chromatin profiling identifies AP1 as a transcriptional regulator in oesophageal adenocarcinoma. PLoS Genet. 13, e1006879.

Burke, Z.D., and Tosh, D. (2012) Barrett’s metaplasia as a paradigm for understanding the development of cancer. Curr Opin Genet Dev. 22:494–9

Chitforoushzadeh Z, Ye Z, Sheng Z, LaRue S, Fry RC, Lauffenburger DA, and Janes KA. (2016) TNF-insulin crosstalk at the transcription factor GATA6 is revealed by a model that links signaling and transcriptomic data tensors. Sci Signal. 9:ra59.

Chia, N.-Y., Deng, N., Das, K., Huang, D., Hu, L., Zhu, Y., Lim, K.H., Lee, M.-H., Wu, J., Sam, X.X., et al. (2015). Regulatory crosstalk between lineage-survival oncogenes KLF5, GATA4 and GATA6 cooperatively promotes gastric cancer development. Gut 64, 707–719.

Colleypriest, B.J., Burke, Z.D., Griffiths, L.P., Chen, Y., Yu, W.Y., Jover, R., Bock, M., Biddlestone, L., Quinlan, J.M., Ward, S.G., et al. (2017). Hnf4α is a key gene that can generate columnar metaplasia in oesophageal epithelium. Differentiation 93, 39–49.

Cusanovich, D.A., Reddington, J.P., Garfield, D.A., Daza, R.M., Aghamirzaie, D., Marco-Ferreres, R., Pliner, H.A., Christiansen, L., Qiu, X., Steemers, F.J., et al. (2018). The cis-regulatory dynamics of embryonic development at single-cell resolution. Nature 555, 538–542.

Davie, K., Jacobs, J., Atkins, M., Potier, D., Christiaens, V., Halder, G., and Aerts, S. (2015). Discovery of Transcription Factors and Regulatory Regions Driving In Vivo Tumor Development by ATAC-seq and FAIRE-seq Open Chromatin Profiling. PLoS Genet. 11, 1–24.

Denny, S.K., Yang, D., Chuang, C.H., Brady, J.J., Lim, J.S.S., Grüner, B.M., Chiou, S.H., Schep, A.N., Baral, J., Hamard, C., et al. (2016). Nfib Promotes Metastasis through a Widespread Increase in Chromatin Accessibility. Cell 166, 328–342.

Desai, T.K., Krishnan, K., Samala, N., Singh, J., Cluley, J., Perla, S., and Howden, C.W. (2012). The incidence of oesophageal adenocarcinoma in non-dysplastic Barrett’s oesophagus: A meta-analysis. Gut 61, 970–976.

Dobin A, Davis CA, Schlesinger F, Drenkow J, Zaleski C, Jha S, Batut P, Chaisson M, Gingeras TR. (2013) STAR: ultrafast universal RNA-seq aligner. Bioinformatics. 29, 15–21.

Dulak, A.M., Stojanov, P., Peng, S., Lawrence, M.S., Fox, C., Stewart, C., Bandla, S., Imamura, Y., Schumacher, S.E., Shefler, E., et al. (2013). Exome and whole-genome sequencing of esophageal adenocarcinoma identifies recurrent driver events and mutational complexity. Nat. Genet. 45, 478–486.

Fitzgerald, R.C., Di Pietro, M., Ragunath, K., Ang, Y., Kang, J.Y., Watson, P., Trudgill, N., Patel, P., Kaye, P. V., Sanders, S., et al. (2014). British Society of Gastroenterology guidelines on the diagnosis and management of Barrett’s oesophagus. Gut 63, 7–42.

Frankell, A.M., Jammula, S., Contino, G., Killcoyne, S.S., Abbas, S., Perner, J., Bower, L., Devonshire, G., Grehan, N., Mok, J., O’Donovan, M., Macrae, S., Tavare, S., Fitzgerald, R.C., the Oesophageal Cancer Clinical and Molecular Stratification (OCCAMS) Consortium. (2018) The landscape of selection in 551 Esophageal Adenocarcinomas defines genomic biomarkers for the clinic. BioRkiv. doi: https://doi.org/10.1101/310029.

Garcia E, Hayden A, Birts C, Britton E, Cowie A, Pickard K, Mellone M, Choh C, Derouet M, Duriez P, Noble F, White MJ, Primrose JN, Strefford JC, Rose-Zerilli M, Thomas GJ, Ang Y, Sharrocks AD, Fitzgerald RC, Underwood TJ; OCCAMS consortium. (2016) Authentication and characterisation of a new oesophageal adenocarcinoma cell line: MFD-1. Sci Rep. 6:32417.

Gerrard DT, Berry AA, Jennings RE, Piper Hanley K, Bobola N, Hanley NA. (2016) An integrative transcriptomic atlas of organogenesis in human embryos. Elife. 5, e15657.

Green, N.H., Nicholls, Z., Heath, P.R., Cooper-Knock, J., Corfe, B.M., Macneil, S., and Bury, J.P. (2014). Pulsatile exposure to simulated reflux leads to changes in gene expression in a 3D model of oesophageal mucosa. Int. J. Exp. Pathol. 95, 216–228.

Gosalia, N., Yang, R., Kerschner, J.L., and Harris, A. (2015). FOXA2 regulates a network of genes involved in critical functions of human intestinal epithelial cells. Physiol. Genomics 47, 290–297.

Heinz, S., Benner, C., Spann, N., Bertolino, E., Lin, Y.C., Laslo, P., Cheng, J.X., Murre, C., Singh, H., and Glass, C.K. (2010). Simple Combinations of Lineage-Determining Transcription Factors Prime cis-Regulatory Elements Required for Macrophage and B Cell Identities. Mol. Cell 38, 576–589.

Jiang, M., Li, H., Zhang, Y., Yang, Y., Lu, R., Liu, K., Lin, S., Lan, X., Wang, H., Wu, H., et al. (2017). Transitional basal cells at the squamous-columnar junction generate Barrett’s oesophagus. Nature 550, 529–533.

Kerschner, J.L., Gosalia, N., Leir, S.-H., and Harris, A. (2014). Chromatin remodeling mediated by the FOXA1/A2 transcription factors activates CFTR expression in intestinal epithelial cells. Epigenetics 9, 557–565.

Langmead, B., Trapnell, C., Pop, M., and Salzberg, S. (2009). Ultrafast and memory-efficient alignment of short DNA sequences to the human genome. Genome Biol. 10, R25.

Lin, L., Bass, A.J., Lockwood, W.W., Wang, Z., Silvers, A.L., Thomas, D.G., Chang, A.C., Lin, J., Orringer, M.B., Li, W., et al. (2012). Activation of GATA binding protein 6 (GATA6) sustains oncogenic lineage-survival in esophageal adenocarcinoma. Proc. Natl. Acad. Sci. 109, 4251–4256.

Maag, J., Fisher, O.M., Levert-Mignon, A.J., Kaczorowski, D.C., Thomas, M.L., Hussey, D., Watson, D., Wettstein, A., Bobryshev, Y. V., Edwards, M., et al. (2017). Novel Aberrations Uncovered in Barrett’s Esophagus and Esophageal Adenocarcinoma Using Whole Transcriptome Sequencing. Mol. Cancer Res. 15:1558–1569.

Mallanna, S.K., Cayo, M.A., Twaroski, K., Gundry, R.L., and Duncan, S.A. (2016). Mapping the Cell-Surface N-Glycoproteome of Human Hepatocytes Reveals Markers for Selecting a Homogeneous Population of iPSC-Derived Hepatocytes. Stem Cell Reports 7, 543–556.

Nowicki-Osuch, K., Li, Y., Challinor, M., Gerrard, D.T., Hanley, N.A., and Sharrocks, A.D. (2017). EINCR1 is an EGF inducible lincRNA overexpressed in lung adenocarcinomas. PLoS ONE 12, e0181902.

Pastushenko, I., Brisebarre, A., Sifrim, A., Fioramonti, M., Revenco, T., Boumahdi, S., Van Keymeulen, A., Brown, D., Moers, V., Lemaire, S., et al. (2018). Identification of the tumour transition states occurring during EMT. Nature 556, 463–468.

Pennathur, A., Gibson, M.K., Jobe, B.A., and Luketich, J.D. (2013). Oesophageal carcinoma. Lancet 381, 400–12.

Piessen, G., Jonckheere, N., Vincent, A., Hemon, B., Ducourouble, M.P., Copin, M.C., Mariette, C., and Van Seuningen, I. (2007). Regulation of the human mucin MUC4 by taurodeoxycholic and taurochenodeoxycholic bile acids in oesophageal cancer cells is mediated by hepatocyte nuclear factor 1alpha. Biochem J 402, 81–91.

Piper, J., Elze, M.C., Cauchy, P., Cockerill, P.N., Bonifer, C., and Ott, S. (2013). Wellington: A novel method for the accurate identification of digital genomic footprints from DNase-seq data. Nucleic Acids Res. 42, 11272.

Quante, M., Bhagat, G., Abrams, J.A., Marache, F., Good, P., Lee, M.D., Lee, Y., Friedman, R., Asfaha, S., Dubeykovskaya, Z., et al. (2012). Bile acid and inflammation activate gastric cardia stem cells in a mouse model of barrett-like metaplasia. Cancer Cell 21, 36–51.

Quinlan, A.R., and Hall, I.M. (2010). BEDTools: A flexible suite of utilities for comparing genomic features. Bioinformatics 26, 841–842.

Ramírez, F., Dündar, F., Diehl, S., Grüning, B.A., and Manke, T. (2014) deepTools: a flexible platform for exploring deep-sequencing data. Nucleic Acids Res. 42, W187–91.

Rendeiro, A.F., Schmidl, C., Strefford, J.C., Walewska, R., Davis, Z., Farlik, M., Oscier, D., and Bock, C. (2016). Chromatin accessibility maps of chronic lymphocytic leukaemia identify subtype-specific epigenome signatures and transcription regulatory networks. Nat. Commun. 7, 11938.

Ross-Innes, C.S., Becq, J., Warren, A., Cheetham, R.K., Northen, H., O’Donovan, M., Malhotra, S., Di Pietro, M., Ivakhno, S., He, M., et al. (2015). Whole-genome sequencing provides new insights into the clonal architecture of Barrett’s esophagus and esophageal adenocarcinoma. Nat. Genet. 47, 1038–1046.

San Roman, A.K., Aronson, B.E., Krasinski, S.D., Shivdasani, R.A., and Verzi, M.P. (2015). Transcription factors GATA4 and HNF4A control distinct aspects of intestinal homeostasis in conjunction with transcription factor CDX2. J. Biol. Chem. 290, 1850–1860.

Sherwood, R.I., Chen, T.Y.A., and Melton, D.A. (2009). Transcriptional dynamics of endodermal organ formation. Dev. Dyn. 238, 29–42.

Spechler, S.J., and Souza, R.F. (2014). Barrett’s Esophagus. N. Engl. J. Med. 371, 836–845.

Stachler, M.D., Taylor-Weiner, A., Peng, S., McKenna, A., Agoston, A.T., Odze, R.D., Davison, J.M., Nason, K.S., Loda, M., Leshchiner, I., et al. (2015). Paired exome analysis of Barrett’s esophagus and adenocarcinoma. Nat. Genet. 47, 1047–1055.

Stairs, D.B., Nakagawa, H., Klein-Szanto, A., Mitchell, S.D., Silberg, D.G., Tobias, J.W., Lynch, J.P., and Rustgi, A.K. (2008). Cdx1 and c-Myc foster the initiation of transdifferentiation of the normal esophageal squamous epithelium toward Barrett’s esophagus. PLoS One 3, e3534.

Sulahian, R., Casey, F., Shen, J., Qian, Z.R., Shin, H., Ogino, S., Weir, B. a, Vazquez, F., Liu, X.S., Hahn, W.C., et al. (2013). An integrative analysis reveals functional targets of GATA6 transcriptional regulation in gastric cancer. Oncogene 33, 1–12.

The Cancer Genome Atlas Research Network (2017). Integrated genomic characterization of oesophageal carcinoma. Nature 541, 169–175.

Toska, E., Osmanbeyoglu, H.U., Castel, P., Chan, C., Hendrickson, R.C., Elkabets, M., Dickler, M.N., Scaltriti, M., Leslie, C.S., Armstrong, S.A., and Baselga, J. (2017) PI3K pathway regulates ER-dependent transcription in breast cancer through the epigenetic regulator KMT2D. Science 355, 1324–1330.

Trapnell, C., Roberts, A., Goff, L., Pertea, G., Kim, D., Kelley, D.R., Pimentel, H., Salzberg, S.L., Rinn, J.L., and Pachter, L. (2012). Differential gene and transcript expression analysis of RNA-seq experiments with TopHat and Cufflinks. Nat. Protoc. 7, 562–578.

Tripathi, S., Pohl, MO., Zhou, Y., Rodriguez-Frandsen, A., Wang, G., Stein, DA., et al., (2015) Meta-and Orthogonal Integration of Influenza "OMICs" Data Defines a Role for UBR4 in Virus Budding. Cell Host Microbe. 18, 723–35.

Vega, M.E., Giroux, V., Natsuizaka, M., Liu, M., Klein-Szanto, A.J., Stairs, D.B., Nakagawa, H., Wang, K.K., Wang, T.C., Lynch, J.P., et al. (2014). Inhibition of notch signaling enhances transdifferentiation of the esophageal squamous epithelium towards a Barrett’s-like metaplasia via KLF4. Cell Cycle 13, 3857–3866.

Verzi, M.P., Shin, H., He, H.H., Sulahian, R., Meyer, C.A., Montgomery, R.K., Fleet, J.C., Brown, M., Liu, X.S., and Shivdasani, R.A. (2010). Differentiation-Specific Histone Modifications Reveal Dynamic Chromatin Interactions and Partners for the Intestinal Transcription Factor CDX2. Dev. Cell 19, 713–726.

Vuong LM, Chellappa K, Dhahbi JM, Deans JR, Fang B, Bolotin E, Titova NV, Hoverter NP, Spindler SR, Waterman ML, and Sladek FM. (2015) Differential Effects of Hepatocyte Nuclear Factor 4a Isoforms on Tumor Growth and T-Cell Factor 4/AP-1 Interactions in Human Colorectal Cancer Cells. Mol Cell Biol. 35:3471–90.

Walker, E.M., Thompson, C.A., and Battle, M.A. (2014). GATA4 and GATA6 regulate intestinal epithelial cytodifferentiation during development. Dev. Biol. 392, 283–294.

Wapinski, O.L., Lee, Q.Y., Chen, A.C., Li, R., Corces, M.R., Ang, C.E., Treutlein, B., Xiang, C., Baubet, V., Suchy, F.P., et al. (2017). Rapid Chromatin Switch in the Direct Reprogramming of Fibroblasts to Neurons. Cell Rep. 20, 3236–3247.

Weaver, J.M.J., Ross-Innes, C.S., Shannon, N., Lynch, A.G., Forshew, T., Barbera, M., Murtaza, M., Ong, C.A.J., Lao-Sirieix, P., Dunning, M.J., et al. (2014). Ordering of mutations in preinvasive disease stages of esophageal carcinogenesis. Nat. Genet. 46, 837–843.

Wei, T. and Simko V. (2017). R package "corrplot": Visualization of a Correlation Matrix (Version 0.84). Available from https://github.com/taiyun/corrplot

Wiseman, E.F., Chen, X., Han, N., Webber, A., Ji, Z., Sharrocks, A.D., and Ang, Y.S. (2015) Deregulation of the FOXM1 target gene network and its coregulatory partners in oesophageal adenocarcinoma. Mol Cancer. 14, 69.

Yamamoto Y, Wang X, Bertrand D, Kern F, Zhang T, Duleba M, Srivastava S, Khor CC, Hu Y, Wilson LH, Blaszyk H, Rolshud D, Teh M, Liu J, Howitt BE, Vincent M, Crum CP, Nagarajan N, Ho KY, McKeon F, Xian W. (2016) Mutational spectrum of Barrett’s stem cells suggests paths to initiation of a precancerous lesion. Nat Commun. 7, 10380.

Yang, R., Kerschner, J.L., and Harris, A. (2016). Hepatocyte nuclear factor 1 coordinates multiple processes in a model of intestinal epithelial cell function. Biochim. Biophys. Acta 1859, 591–598.

Yu, V.W.C., Yusuf, R.Z., Oki, T., Wu, J., Saez, B., Wang, X., Cook, C., Baryawno, N., Ziller, M.J., Lee, E., et al. (2016). Epigenetic memory underlies cell-autonomous heterogeneous behavior of hematopoietic stem cells. Cell 167, 1310–1322.e17.

Zaret KS, and Carroll JS. (2011) Pioneer transcription factors: establishing competence for gene expression. Genes Dev. 25:2227–41.

Zhang, Y., Liu, T., Meyer, C.A., Eeckhoute, J., Johnson, D.S., Bernstein, B.E., Nussbaum, C., Myers, R.M., Brown, M., Li, W., et al. (2008). Model-based analysis of ChIP-Seq (MACS). Genome Biol. 9, R137.

